# Cellular Immunotherapy in Mice Prevents Maternal Hypertension and Restores Anti-Inflammatory Cytokine Balance in Maternal and Fetal Tissues

**DOI:** 10.1101/2023.08.14.553300

**Authors:** Gabrielle Gray, Douglas G. Scroggins, Katlin T. Wilson, Sabrina M. Scroggins

**Affiliations:** Department of Biomedical Sciences, University of Minnesota Duluth, Duluth, MN 55812 USA; University of Iowa, Iowa City, IA 52242 USA

**Keywords:** Cellular Therapy, Preeclampsia, Cytokines, Inflammation, Regulatory Dendritic Cells

## Abstract

Preeclampsia is the leading cause of maternal-fetal morbidity world-wide.

The concept that persistent feto-placental intolerance is important in the pathogenesis of preeclampsia (PreE) has been demonstrated by our lab and others. Arginine vasopressin (AVP) infusion during pregnancy induces cardiovascular, renal, and T helper (Th) cell alterations in mice consistent with human PreE. In addition to their conventional immuno-stimulatory role, dendritic cells (DCs) also play a vital role in immune tolerance. In contrast to conventional DCs, regulatory DCs (DCregs) express low levels of co-stimulatory markers, produce anti-inflammatory cytokines, induce T regulatory (Treg) cells, and promote tolerance. In mice, DCregs prevent pro-inflammatory responses and induce antigen-specific tolerance. Given these known functions of DCregs, we hypothesize that DCregs will prevent the development of AVP-induced PreE in mice. C57BL/6J females were infused with AVP (24 ng/hour) or saline throughout gestation via osmotic minipump. Bone marrow derived DCregs were injected into AVP-infused dams at the time of pump implantation or on gestational day (GD) 7. Blood pressure was taken throughout pregnancy. Maternal urine protein and TH-associated cytokines in maternal and fetal tissues were measured on GD 18. Treatment with DCregs effectively prevented the elevation of maternal blood pressure, proteinuria, and fetal growth restriction that was observed in AVP-infused dams. Furthermore, we noted a reduction in pro-inflammatory TH-associated cytokines IFNγ and IL-17, while anti-inflammatory cytokines IL-4, IL-10, and TGFβ showed an increase following DCreg treatment. These outcomes provide strong evidence supporting the potential of DCregs as a valuable therapeutic approach in addressing PreE.

## 1. Introduction

Preeclampsia (PreE), a hypertensive disorder in pregnancy, affects 5-7% of all pregnancies in the United States, equating to 400,000 pregnancies annually. It is a leading cause of maternal-fetal mortality worldwide resulting in approximately 76,000 maternal and 500,000 neonatal deaths each year. PreE causes both immediate and long-term medical complications for the fetus and mother [1]. Mothers affected by PreE and their children are at increased risk of developing cardiovascular and metabolic disease as well as cognitive impairments later in life [2-14]. Currently, the only curative option is the often-preterm delivery of the fetus, which contributes considerable fetal morbidity and mortality [15], underscoring the importance of developing new modalities to prevent and treat PreE.

The pathogenesis of PreE involves multiple bodily systems, including placental, vascular, renal, and immune dysfunction. Because PreE is a disease resulting from these multiple pathways, successful interventions to prevent and treat PreE must originate from the upstream initiation and regulation of these multiple pathways. Our previous work and others demonstrating T helper (TH) cell dysregulation contributes to poor vascular, renal, and immune function in PreE [16-21], supports the concept that immune dysfunction, particularly Th cell dysregulation, may play an etiological role in PreE. Modulation of T cells may be an effective treatment strategy for PreE. Dendritic cells (DCs) are antigen presenting cells that play a pivotal role in T cell activation and regulation [22-24]. In addition to their conventional immuno-stimulatory role in adaptive immunity, DCs also play a pivotal role in immune homeostasis and tolerance through direct interactions with T cells and cytokine production [22]. In contrast to conventional DCs, regulatory DCs (DCreg) are defined by their ability to induce T cell tolerance *in vitro* and *in* vivo via direct or indirect induction of Treg and/or T cell anergy [25-30]. In animal models, DCreg prevent lethal systemic inflammatory responses, inflammatory bowel disease, allergic airway disease, solid organ allograft rejection, and lethal graft vs. host disease [31-36] and prove an effective therapy for the treatment of allergic airway and autoimmune diseases [37]. Like PreE, immune dysfunction is pathological in these diseases. Therefore, the overall objective of this study was to determine if administration of DCreg would prevent hypertension and restore immune balance in PreE. To this end, a pregnancy-specific vasopressin (AVP) -induced mouse model of PreE was utilized to assess hallmark features of PreE: elevated blood pressure, proteinuria, fetal growth restriction, and maternal and fetal tissue cytokine profiles on gestational day 18 (GD18).

## 2. Results

### 2.1. DCreg Administration Prevents Maternal and Fetal PreE-Associated Morbidities

Blood pressure was taken throughout gestation via tail cuff on saline or AVP-infused dams. As previously published in rodent models [21, 38-40], AVP infusion during pregnancy resulted in significantly higher systolic blood pressure (Figure 1A), increased 24-hour urine protein concentration (Figure 1B), and lower pup weights (Figure 1E) with no remarkable impact on the size of litters (Figure 1C) or resorptions per litter (Figure 1D). Administration of DCreg at the time of mini-pump implantation (GD-3) or on GD7 prevented increases in systolic blood pressure (Figure 1A), urine protein (Figure 1B), and growth restriction (Figure 1E). DCreg administration did not alter the number of pups (Figure 1C) or resorptions (Figure 1D) per litter. Collectively, these data support DCreg administration in the prevention of AVP-induced increases in blood pressure, proteinuria, and growth restriction.

**Figure 1.**
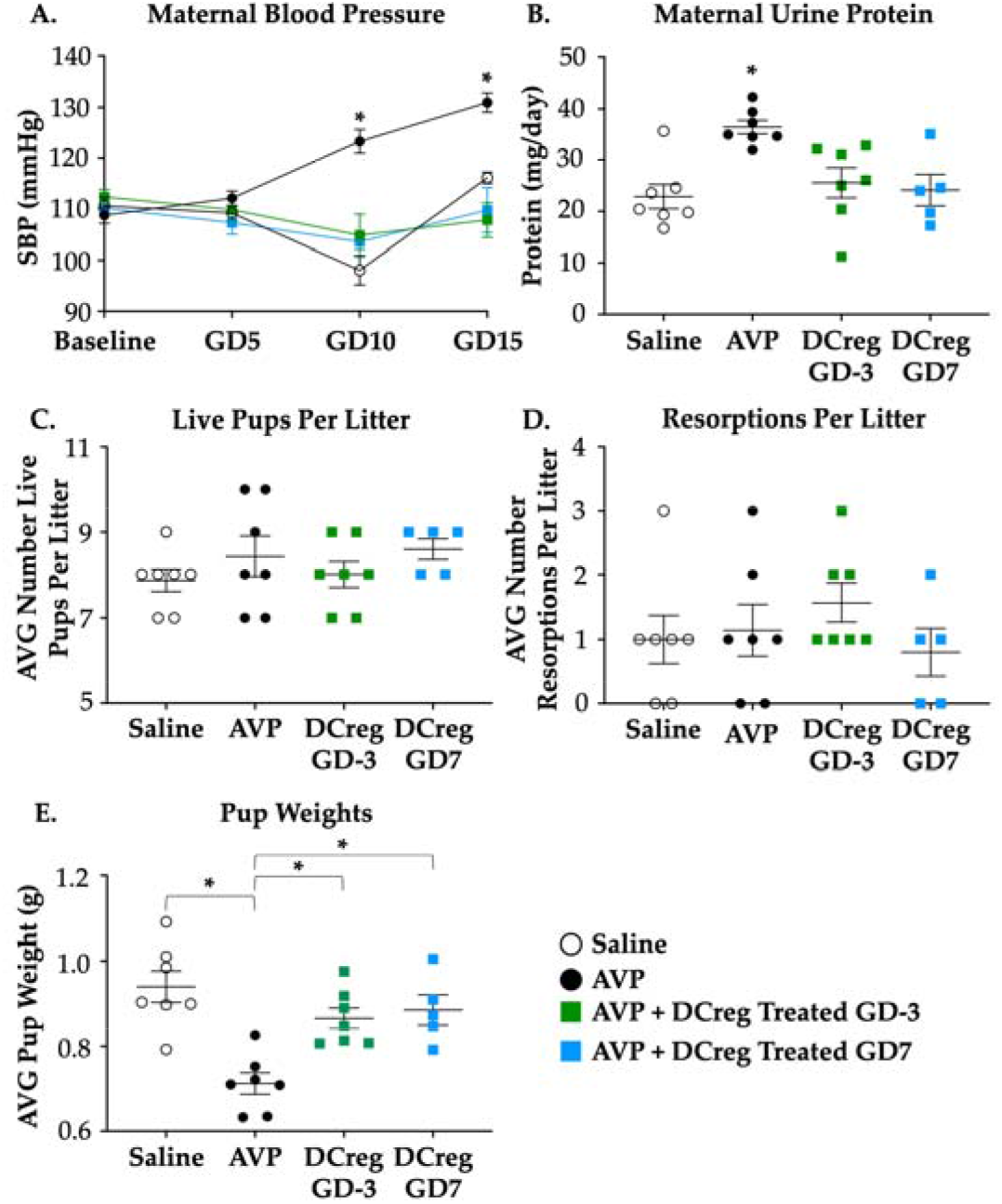
DCreg administration prevents increases in blood pressure and proteinuria in vasopressin-induced preeclampsia. (**a**) Systolic blood pressure throughout gestation in vasopressin-induced preeclampsia; (**b**) Maternal 24-hour urine protein concentrations on GD18; (**c**) Average number of live pups per litter; (**d**) Average number of resorbed pups per litter; (**e**) Average weight of pups in each litter at the time of delivery. AVP= vasopressin; GD= gestational day. Open circles= saline; Filled circles= AVP; Green Filled Squares= AVP + DCreg GD-3; Blue Filled Squares= AVP + DCreg GD7. N=5-7 per group. Data are ± SEM. * p<0.05.

### 2.2. DCreg Treatment Prevents Elevated Maternal and Fetal Inflammatory Cytokines

Consistent with previous studies, IFN⍰ (Figure 2A) and IL-17 (Figure 2B) were increased in the serum of AVP-infused dams compared to saline. DCreg administration significantly reduced IFN⍰ and IL-17 in maternal serum at both GD-3 and GD7 with no significant changes in maternal kidney IFN⍰ or IL-17 (Figure 2A and 2B, respectively). Administration of DCreg at either GD-3 or GD7 returned maternal serum IFN⍰ (Figure 2A) and IL-17 (Figure 2B) to concentrations comparable to saline-infused dams. AVP infusion during gestation did not significantly alter IFN⍰ concentrations in the amniotic fluid, placenta, fetal kidney, or fetal liver; and DCreg administration did not alter IFN⍰ concentrations in these fetal tissues (Figure 2C). Maternal AVP-infusion resulted in increased IL-17 in the amniotic fluid, placenta, and fetal kidney (Figure 2D). Maternal DCreg treatment prevented increases in the concentration of IL-17 in the amniotic fluid, placenta, and fetal kidney. Fetal liver IL-17 concentrations were not significantly impacted by maternal AVP-infusion or DCreg administration (Figure 2D).

**Figure 2.**
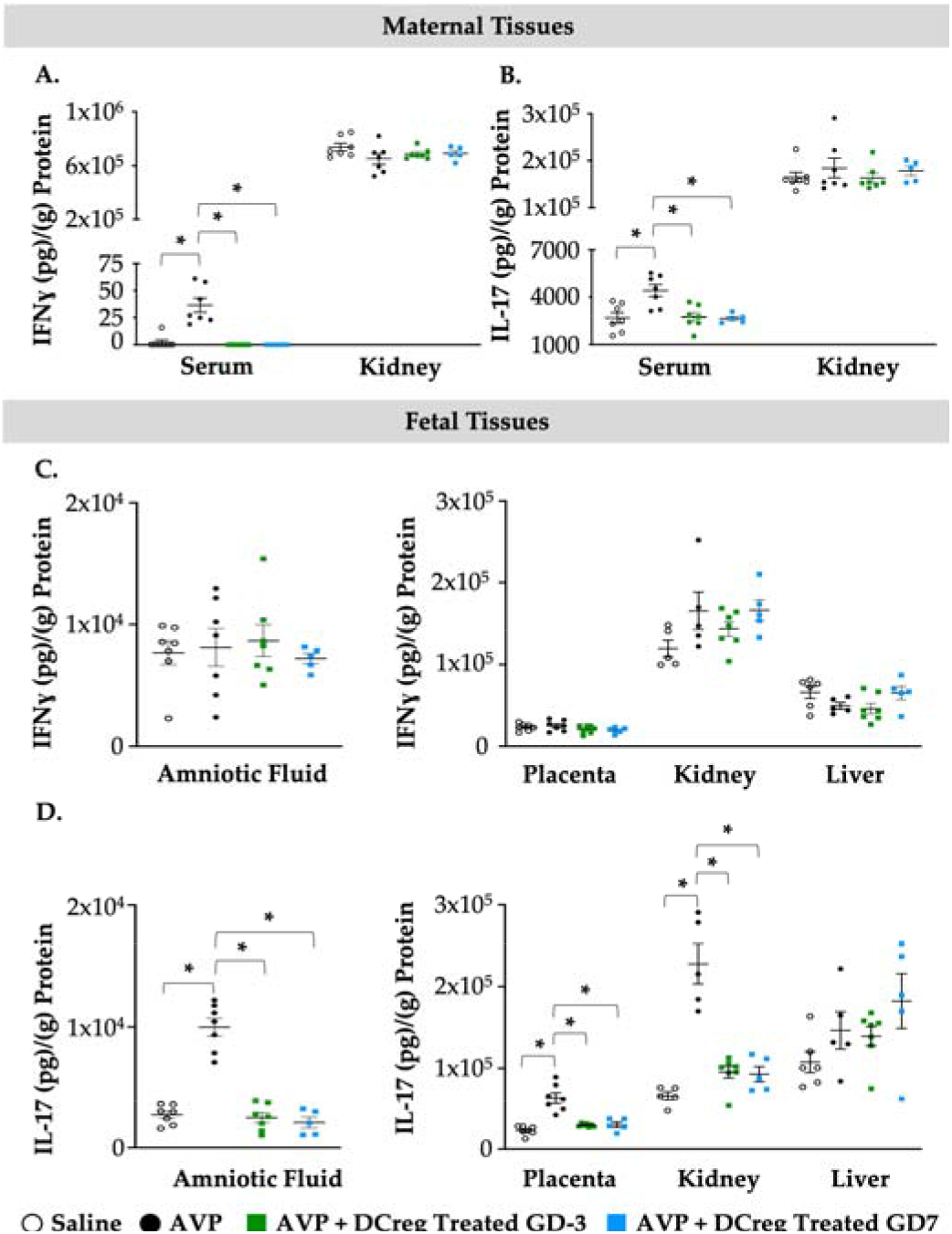
Preeclampsia-induced inflammation in maternal and fetal tissues is ameliorated by maternal DCreg administration. (**a**) IFNl7l and (**b**) IL-17 concentrations in maternal serum and kidney. Concentrations of (**c**) IFNl7l and (**d**) IL-17 in amniotic fluid, placenta, fetal kidney, and fetal liver. AVP= vasopressin; GD= gestational day. Open circles= saline; Filled circles= AVP; Green Filled Squares= AVP + DCreg GD-3; Blue Filled Squares= AVP + DCreg GD7. N=5-7 per group. Data are ± SEM. * p<0.05.

### 2.3. Treatment with DCreg Restores Maternal and Fetal Tissue Anti-Inflammatory Cytokines

AVP-infusion during gestation resulted in decreased anti-inflammatory cytokines IL-4, IL-10, and TGF*β* in the maternal serum and kidney. DCreg treatment prior to mating (GD-3) or on GD7 significantly increased IL-4, IL-10, and TGF*β* within these tissues (Figure 3A-3C). Amniotic fluid, placenta, and fetal kidney IL-4 and IL-10 were significantly decreased in offspring from AVP-infused dams (Figure 3D, 3E, 3G, and 3H), while TGF*β* was significantly decreased in the placenta and fetal kidney (Figure 3F and 3I). Treatment of AVP-infused dams on GD-3 or GD7 prevented AVP-induced suppression of IL-4, IL-10, and TGF*β* in the amniotic fluid, placenta, and fetal kidney (Figure 3D-I). In the fetal liver, only IL-10 was signficantly reduced due to AVP-infusion and DCreg treatment at either timepoint restored IL-10 levels (Figure 3H).

**Figure 3.**
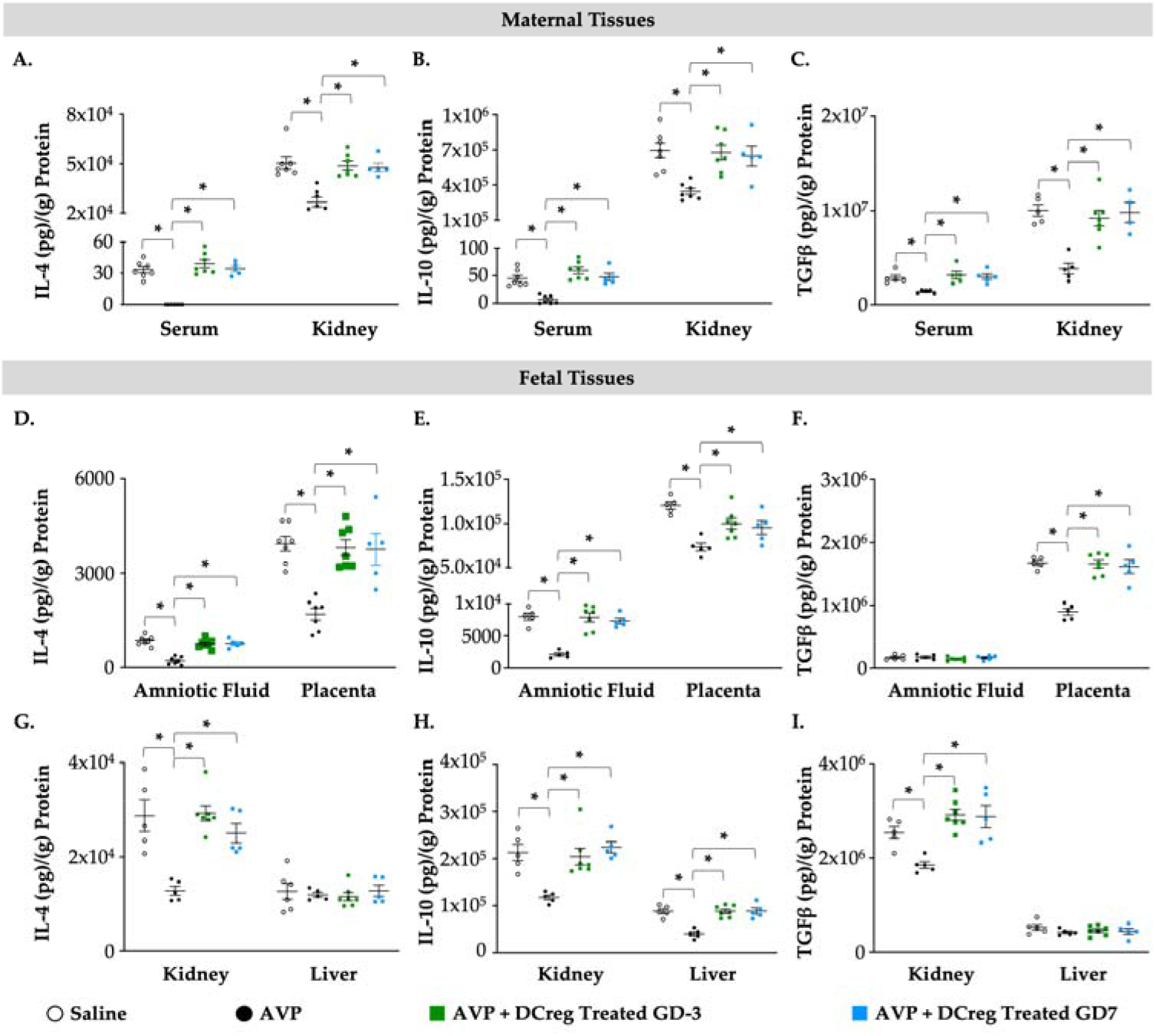
Administration of DCreg upregulates production of anti-inflammatory cytokines in maternal and fetal tissues from preeclampsia-affected pregnancies. Concentrations of (**a**) IL-4, (**b**) IL-10, and (**c**) TGF*β* in maternal serum and kidney. Amniotic fluid and placenta concentrations of (**d**) IL-4, (**e**) IL-10, and (**f**) TGF*β*. Concentrations of (**g**) IL-4, (**h**) IL-10, and (**i**) TGF*β* in fetal kidney and liver at GD18. AVP= vasopressin; GD= gestational day. Open circles= saline; Filled circles= AVP; Green Filled Squares= AVP + DCreg GD-3; Blue Filled Squares= AVP + DCreg GD7. N=5-7 per group. Data are ± SEM. * p<0.05.

## 3. Discussion

PreE is the leading cause of maternal-fetal morbidity in both developed and developing countries. Currently, there is a lack of effective interventions to prevent and treat PreE. Immune dysfunction and enhanced T cell-mediated inflammation are key to the pathogenesis of PreE. Therefore, the objective of the current study was to determine if modulation of immune responses via cellular immunotherapy mitigates key aspects of PreE.

Although PreE begins very early in pregnancy, one of the first symptoms of PreE in humans and mouse models is elevated blood pressure. Later in gestation, proteinuria and fetal growth restriction are associated with PreE. Indeed, our AVP-induced mouse model demonstrates increased blood pressure, proteinuria, and fetal growth restriction. Administration of DCreg prevented the PreE-induced changes in vascular, renal, and fetal outcomes. A likely mechanism of DCreg alleviation of PreE is the regulation of inflammatory T cell responses in PreE.

In instances of PreE, there is an increase in TH1 cells producing IFNγ and TH17 cells producing IL-17, along with a decrease in TH2-related cytokines like IL-4 and IL-10 [18, 21, 41-43]. This creates an environment abundant in both cells and cytokines that encourage an inflammatory response. While the precise mechanisms responsible for initiating PreE are complex and not fully understood, it is clear that an atypical Th cell response to pregnancy has a significant impact [16]. Indeed, we have observed elevated levels of IFNγ and IL-17, alongside reduced levels of IL-4, IL-10, and TGF*β* in both maternal and fetal tissues from pregnancies affected by PreE as well as reduced proportions of Treg cells in PreE [21]. These alterations in TH-associated cytokines were prevented through treatment of dams before and early in the pregnancy with DCreg.

Dendritic cells (DCs) are widely acknowledged as the primary antigen-presenting cells that contribute to adaptive immunity [25, 26]. Beyond their conventional role in immune stimulation, DCs also hold a crucial position in maintaining immune balance and tolerance through their interactions with T cells and cytokine secretion [24]. While conventional DCs have their immuno-stimulatory functions, DCregs induce T cell tolerance both *in vitro* and *in vivo*. This tolerance-inducing process can happen through direct or indirect induction of Tregs and/or T cell anergy [38, 54-58]. DCregs exhibit a distinctive ratio of co-stimulatory to inhibitory surface molecules. Alongside their minimal expression of co-stimulatory molecules, DCregs also generate anti-inflammatory cytokines [20, 38, 59-61].

Various identifying features and functional attributes have been associated with DC populations possessing regulatory capabilities. These include the expression of PD-L1 and PD-L2 [27, 29], as well as the production of IL-10 and TGF-*β* [32-35, 65-68], and indoleamine 2,3 dioxygenase (IDO) [69, 70]. DCregs that produce IL-10 and IDO facilitate the differentiation of Tregs. Previous research has strongly indicated the involvement of DCregs in fostering tolerance towards allo-antigens [33, 71]. In our study, DCreg were able to prevent PreE-induced Th cell-associated dysregulation. A study by Zhang et al. demonstrated that the PD-1/PD-L1 pathway contributes to the imbalance of Treg and TH17 cells in PreE [41]. In a previous study, we showed reduced expression of PD-L1 on DCs from PreE-affected dams [21]. One potential mechanism for the prevention of PreE-induced inflammation, in the current study may be via alteration of the PD-1/PD-L1 pathway. Additional studies are ongoing to investigate the role of the PD-1/PD-L1 and other known regulatory pathways in DCreg-mediated PreE prevention.

## 4. Materials and Methods

### 4.1. Generation of DCreg

Eight days before timed mating, syngeneic DCreg cultures were started as previously described by our group and others [36, 42]. Briefly, bone marrow cells from female C57BL/6J mice were cultured at 1x105 with human TGF-*β*1, and murine GM-CSF and IL-10 (Peprotech; 20 ng/mL each). To terminally mature DCreg before intravenous adoptive transfer, LPS (Sigma-Aldrich; 1 *μ*g/mL) was added to DCreg cultures on day 6 of culture. On day 8 of culture, DCreg were isolated and washed with sterile 0.9% sodium chloride. Cells were resuspended in 0.9% sodium chloride and 5x106 DCregs were administered intravenously per mouse.

### 4.2. Induction of Preeclampsia

Because PreE is a maternal disease that only occurs in females, only females were used to induce preeclampsia in mice. To ensure accurate gestational calculations, all mating in the current study was for a single over-night period (GD 0) using wild-type male C57BL/6J mice. Three days prior to mating (GD -3), 12-week-old virgin female C57BL/6J mice were implanted with a subcutaneous osmotic mini-pumps (Alzet), to deliver 24 ng/hr AVP (Tocris, BioTechne, Minneapolis, MN) or 0.9% sodium chloride (saline; Vetivex, Dechra Veterinary Products, Leawood, KS) [21, 39]. For DCreg-treated groups, DCreg were intravenously administered at the time of pump implantation (GD-3) or on GD7. The four experimental groups of dams were: 1) Saline (control), 2) AVP-infused (preeclamptic), 3) AVP-infused + GD-3 DCreg-treated, and 4) AVP-infused + GD7 DCreg-treated.

On GD18, the dams were euthanized and underwent maternal and fetal necropsy. Litters size and pup weights were recorded at the time of maternal euthanasia. Each pregnancy was designated N=1. Fetal tissues from a single pregnancy were pooled from 5 separate fetuses for an N=1. Data are pooled from multiple independent experiments. Maternal tissues (serum and kidney) and fetal tissues (amniotic fluid, placenta, kidney, and liver) had N≥5 per group from a minimum of two independent experiments. All samples were stored at -80°C for subsequent protein extraction, quantification, and cytokine analysis.

### 4.3. Whole Tissue Extraction of Protein and Total Protein Analysis

Whole maternal and fetal tissues were homogenized in buffer containing 5M NaCl, 1M Tris, 0.5M EDTA, NP-40, protease inhibitor (Roche, Switzerland), and phosphatase inhibitor (Roche, Switzerland). Total protein was quantified using a bicinchoninic acid (BCA) protein assay kit (Thermo Fisher Scientific, Waltham, MA) per manufacturer’s instructions. ELISAs were used to quantify pro-inflammatory IFN⍰ and IL-17 and anti-inflammatory IL-4, IL-10, and TGF*β* of all samples in duplicate (Invitrogen, Thermo Fisher Scientific, Waltham, MA) Cytokine concentrations were normalized to total protein (grams) and are represented as picograms/gram of total protein (pg/g).

### 4.4. Blood Pressure and Proteinuria

Systolic blood pressure (SBP) was obtained as we previously described using the clinically validated CODA Noninvasive Blood Pressure System (Kent Scientific Corp, Torrington, CT) [39, 43, 44]. Mice were acclimated to the CODA system for 21 days prior to pump implantation. Measurements throughout the study were collected at the same time of day, seven days/week. A total of 60 cycles were collected daily. The first 30 cycles each day were considered daily acclimation and discarded. The last 30 cycles each day were used to calculate mean daily measurements for each mouse. The last 5 days of recording during the initial 21-day acclimation period were used to calculate baseline blood pressure prior to pump implantation. Experimental blood pressure data was collected through GD16. On GD16, dams were placed into metabolic cages for 24-hour urine collection. Total protein concentrations of 24-hour GD18 urine was determined using a bicinchoninic acid (BCA) protein assay kit (Thermo Fisher Scientific, Waltham, MA) per manufacturer’s instructions. Urine protein is reported as mg protein 24-hours.

### 4.5. Statistical Analysis

For continuous variables, a two-tailed Student’s t test with unequal variance was utilized, one-way, or two-way ANOVA multiple comparisons test where appropriate (GraphPad, Prism 10, La Jolla, CA). Data are expressed as ± standard error of the mean (SEM). The minimal level of confidence designated statistically significant was α =0.05 or as determined by Bonferroni correction for multiple comparisons ANOVA.

## 5. Conclusions

Our study aimed to determine if DCreg cellular immunotherapy will prevent the cardiovascular, renal, fetal, and immunological alterations associated with PreE. We determined that DCreg treatment prior to pregnancy or early in pregnancy was able to reduce maternal blood pressure, decrease urine protein concentrations, and restore the balance between pro- and antiinflammatory Th cytokines. Furthermore, treatment with DCreg did not increase fetal resorption rates or have any observable differences in fetal outcomes at GD18. Thus, this study establishes that DCreg immunotherapy may be a potent therapeutic intervention for PreE.

## Supplementary Materials

N/A.

## Author Contributions

Conceptualization, S.M.S.; methodology, S.M.S.; experimentation, G.G., D.G.S., and S.M.S; data analysis, G.G. and S.M.S.; writing—original draft preparation, G.G.; writing—review and editing, K.T.W., D.G.S., and S.M.S.; funding acquisition, S.M.S. All authors have read and agreed to the published version of the manuscript.

## Institutional Review Board Statement

All experiments were carried out in accordance with a University of Minnesota Institutional Animal Care and Use Committee approved animal protocol (#2203-39886A approved 07/25/2022).

## Data Availability Statement

The data presented in this study are available upon request from the corresponding author.

## Acknowledgments

The authors would like to acknowledge the unapologetic support from administrative associates Cheryl Anderson and Christine Strom. We would also like to thank Angela Slattery and Patty Sutliff-Opoien for their expertise in grants and financial management.

## Conflicts of Interest

The authors declare no conflict of interest.

### Disclaimer/Publisher’s Note

The statements, opinions and data contained in all publications are solely those of the individual author(s) and contributor(s) and not of MDPI and/or the editor(s). MDPI and/or the editor(s) disclaim responsibility for any injury to people or property resulting from any ideas, methods, instructions or products referred to in the content.

